# A simple RNA preparation method for SARS-CoV-2 detection by RT-qPCR

**DOI:** 10.1101/2020.05.07.083048

**Authors:** Aniela Wozniak, Ariel Cerda, Catalina Ibarra-Henriquez, Valentina Sebastian, Grace Armijo, Liliana Lamig, Carolina Miranda, Marcela Lagos, Sandra Solari, Ana María Guzmán, Teresa Quiroga, Susan Hitschfeld, Eleodoro Riveras, Marcela Ferres, Rodrigo A. Gutiérrez, Patricia García

## Abstract

The technique RT-qPCR for viral RNA detection is the current worldwide strategy used for early detection of the novel coronavirus SARS-CoV-2. RNA extraction is a key pre-analytical step in RT-qPCR, often achieved using commercial kits. However, the magnitude of the COVID-19 pandemic is causing disruptions to the global supply chains used by many diagnostic laboratories to procure the commercial kits required for RNA extraction. Shortage in these essential reagents is even more acute in developing countries with no means to produce kits locally. We sought to find an alternative procedure to replace commercial kits using common reagents found in molecular biology laboratories. Here we report a method for RNA extraction that takes about 40 min to complete ten samples, and is not more laborious than current commercial RNA extraction kits. We demonstrate that this method can be used to process nasopharyngeal swab samples and yields RT-qPCR results comparable to those obtained with commercial kits. Most importantly, this procedure can be easily implemented in any molecular diagnostic laboratory. Frequent testing is crucial for individual patient management as well as for public health decision making in this pandemic. Implementation of this method could maintain crucial testing going despite commercial kit shortages.

## Introduction

SARS-CoV2, a member of the *Coronaviridae* family, is the etiological agent of the current COVID-19 pandemic that has generated an international public health emergency. As of May 3^rd,^ 2020, the virus has infected more than 3.3 million individuals and killed over 238,000 people worldwide (Situation Report 104 of the World Health Organization). Testing for the presence of the virus is of utmost importance for containment strategies aiming to reduce dissemination of the virus and prescription of appropriate clinical practices for affected patients. However, understanding and managing the full extent of the outbreak has remained a challenge for most countries due to significant bottlenecks imposed by diagnosis^1^.

Early detection of infection by SARS-CoV2 relies on the efficient detection of the viral genome using RT-qPCR. Several RT-qPCR-based tests are being used in clinical settings^2^, and novel approaches are constantly being reported^3–10^. All methods require an RNA extraction step to isolate the viral genetic material before its detection. Unfortunately, RNA extraction has become a serious bottleneck for COVID-19 diagnosis around the world due to shortages in RNA-extraction kits customarily used to process patients samples. This is particularly troublesome in developing countries lacking the infrastructure and capacities to produce these kits locally. Before the kit-era, which contributed to standardize and simplify molecular biology work, several RNA extraction methods were routinely used in research laboratories around the world. RNA isolation procedures typically involve three general steps: cell lysis, separation of RNA from other macromolecules such as DNA, proteins, and lipids, followed by RNA concentration. To prevent RNA degradation, cell lysis must be conducted under conditions that inhibit RNase activity, which is abundant in many cellular compartments^11,12^. RNA separation from other macromolecules is often achieved by a combination of pH and organic solvents, such as phenol/chloroform^13–16^. RNA concentration is most commonly achieved by high salt and isopropanol or ethanol precipitation^11,12,17–20^.

We reviewed the published literature to search for procedures of RNA extraction that could potentially be used to replace commercial kits. Many different protocols and variations have been published over the years that optimize or simplify the RNA extraction process from various types of samples. We tested five types of procedures to identify an efficient procedure for extracting RNA from clinical samples that is compatible with downstream RT-qPCR analysis. Of the procedures evaluated, a simple method based in acid pH separation of RNA was found the most suitable. It can be carried out in approximately 40 min for ten samples, and is not more laborious than current methods using commercial kits. This procedure requires reagents and equipment that can be found in any standard molecular biology laboratory, thus avoiding supply chain issues. The resulting RNA can be used to detect SARS-CoV2 by standard RT-qPCR testing protocols with robust results comparable to those obtained using commercial RNA-extraction kits.

## Results

### Screening of alternative procedures for RNA extraction

For validation of the RNA extraction procedures, the RNase P target was amplified in a one-step RT-qPCR reaction, as quality control for the extraction method. As shown in **Figure 1**, three of the five procedures evaluated yielded enough RNA to amplify the target gene, whereas two of them did not. The TRIzol approach was most effective, exhibiting the highest yield when amplifying the human RNase P target **(Figure 1)**. The BSA-based protocol also allowed for amplification of the RNase P target, albeit with a lower yield and significant variability among replicates **(Figure 1)**. Acid pH-based method also allowed amplification of the RNase P target, though with lower yields when compared to the TRIzol method **(Figure 1)**. The direct method and high-temperature method did not yield enough RNA to amplify the RNase P gene under our experimental conditions. While TRIzol appears to be the best experimental procedure in terms of yield, it is not easy to use for a diagnostics laboratory setting as it requires a chemical hood for the organic extraction step. Biosafety cabinets class II (BSL-2) necessary for operator protection are not appropriate for working with organic solvents. BSA, TRIzol, and acid pH procedures provided comparable yields, but the acid pH method was more consistent among replicates. Based on these considerations, we decided to validate the acid pH method to extract RNA from clinical samples, using High Pure Viral RNA Kit (Roche) as the gold standard.

**Figure 1.**
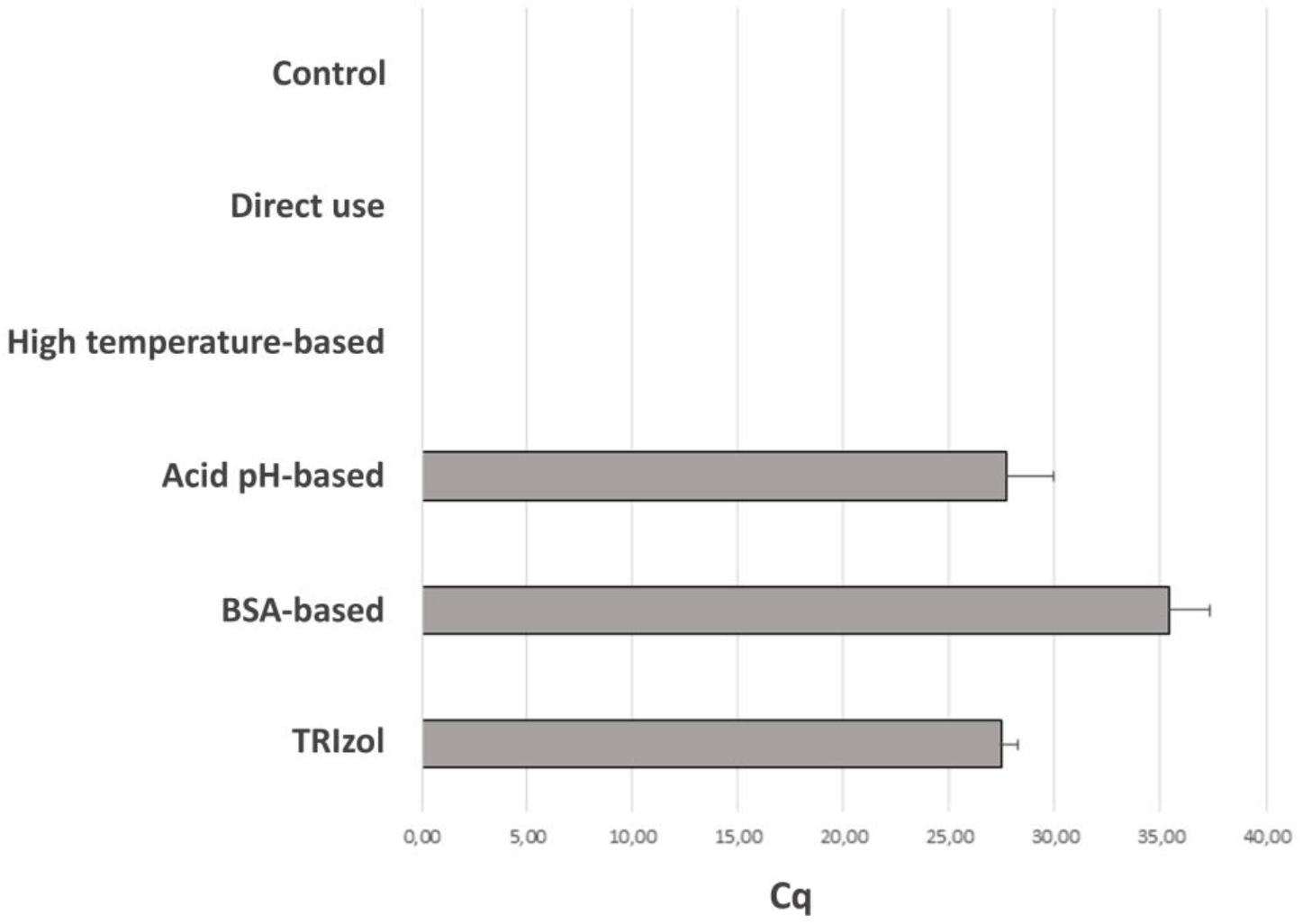
Quantitative assessment of performance for selected RNA extraction methods. Cq values obtained by RT-qPCR with 45 cycles using TaqMan probe and primers against RNase P gene in saliva samples for TRIzol (27.39 +/− 0.34), BSA-based (35.3 +/− 0.79), acid pH-based (27.68 +/− 0.90), high temperature-based (n.d.) and direct (n.d.) methods. n.d.; not determined (no Cq reported). Control corresponds to a negative control with water instead of template. Bars show mean plus standard deviation of the mean for two biological and three technical replicates each (6 measurements).

### Validation of the acid pH RNA extraction method in clinical samples

To validate the acid pH method of RNA extraction, RT-qPCR using TaqMan probes and primers recommended by the CDC were used^21^. The nucleocapsid viral proteins N1 and N2 were amplified as viral targets, and RNase P was also amplified as a control. We analyzed 50 clinical samples: 22 were positive, 11 were undetermined, and 17 were negative according to RT-qPCR recommended by CDC^21^, using RNA extracted with High Pure Viral RNA Kit (Roche). Undetermined samples are described as having a viral load around the limit of detection (LOD) of the RT-qPCR method that was reported as 10^0.5^ RNA copies / μl^21^. This means that the RT-qPCR method can detect 16 RNA copies per PCR reaction. The PCR test used detects 2 targets of the virus: N1 and N2. The mean Cq value for N1 target reported for sets of dilutions that are ≥ 95% positive is around 36^21^. Therefore we analyzed the efficiency of both extraction methods in two different groups of samples: those with Cq N1 ≤36 and those with Cq N1 >36 The results for the 50 samples are shown in **Table 1**. For samples with Cq N1 ≤36 there were no differences in Cq values for N1 and N2 obtained using High Pure Viral RNA Kit (Roche) or the acid pH method **(Figure 2A)**. In contrast, for samples with Cq N1 >36, Cq values for N1 and N2 were higher for High Pure Viral RNA Kit (Roche) than those obtained with acid pH method (p=0.026 and p=0.022 respectively) (**Figure 2B**). For samples with Cq N1 ≤36 Cq values for RNase P were slightly higher for acid pH method (p=0.021), whereas for samples with Cq N1 >36 there was no significant difference between both methods. The % of agreement between both methods was calculated considering samples whose report changed from positive, undetermined or negative. In total, 8 samples changed their report. The 17 negative samples were also negative using RNA extracted with the acid pH method. Out of 22 positive samples, 21 were also positive using RNA extracted with the acid pH method, whereas one sample was undetermined. Out of 11 undetermined samples analyzed, 4 were still undetermined using RNA extracted with the acid pH method. However, 3 of them were negative and 4 of them were positive. Agreement for negative samples was 100%. The percentage of agreement for samples with Cq N1 ≤36 was 89.5% As expected, the percentage of agreement for samples with Cq N1 >36 was only 57%.

**Table 1.**
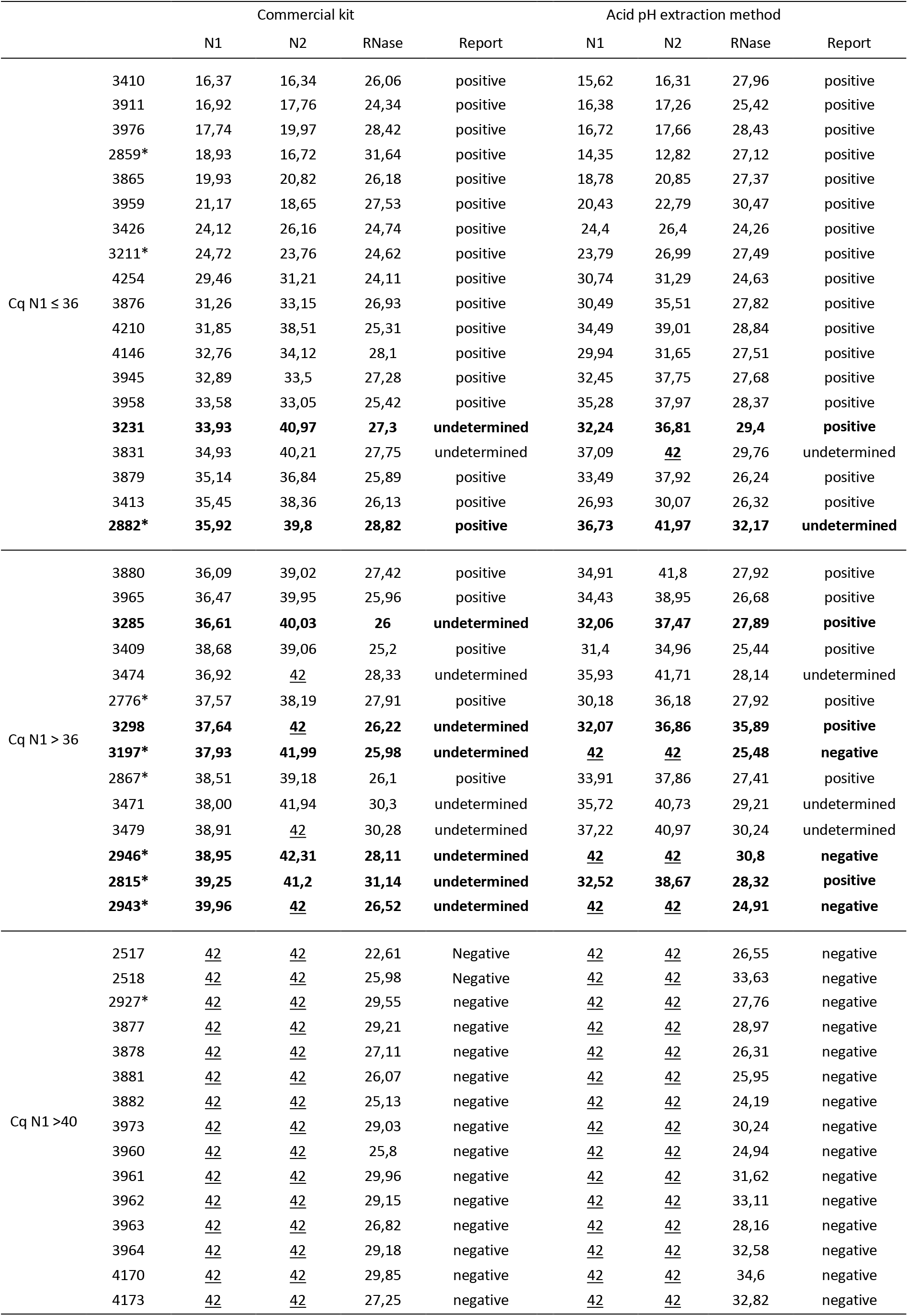
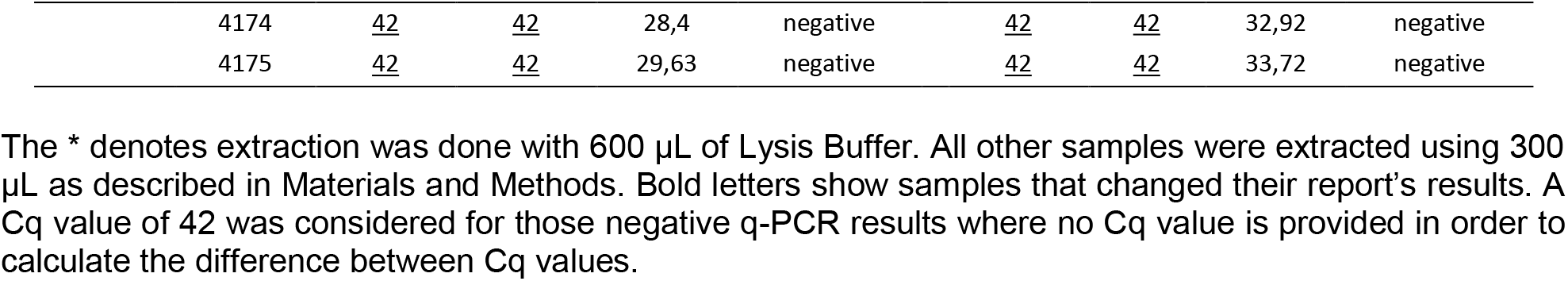
Comparative Cq data for the two RNA extraction methods tested.

|  |  | Commercial kit |  |  |  | Acid pH extraction method |  |  |  |
| --- | --- | --- | --- | --- | --- | --- | --- | --- | --- |
|  |  | N1 | N2 | RNase | Report | N1 | N2 | RNase | Report |
| Cq N1 ≤ 36 | 3410 | 16,37 | 16,34 | 26,06 | positive | 15,62 | 16,31 | 27,96 | positive |
|  | 3911 | 16,92 | 17,76 | 24,34 | positive | 16,38 | 17,26 | 25,42 | positive |
|  | 3976 | 17,74 | 19,97 | 28,42 | positive | 16,72 | 17,66 | 28,43 | positive |
|  | 2859* | 18,93 | 16,72 | 31,64 | positive | 14,35 | 12,82 | 27,12 | positive |
|  | 3865 | 19,93 | 20,82 | 26,18 | positive | 18,78 | 20,85 | 27,37 | positive |
|  | 3959 | 21,17 | 18,65 | 27,53 | positive | 20,43 | 22,79 | 30,47 | positive |
|  | 3426 | 24,12 | 26,16 | 24,74 | positive | 24,4 | 26,4 | 24,26 | positive |
|  | 3211* | 24,72 | 23,76 | 24,62 | positive | 23,79 | 26,99 | 27,49 | positive |
|  | 4254 | 29,46 | 31,21 | 24,11 | positive | 30,74 | 31,29 | 24,63 | positive |
|  | 3876 | 31,26 | 33,15 | 26,93 | positive | 30,49 | 35,51 | 27,82 | positive |
|  | 4210 | 31,85 | 38,51 | 25,31 | positive | 34,49 | 39,01 | 28,84 | positive |
|  | 4146 | 32,76 | 34,12 | 28,1 | positive | 29,94 | 31,65 | 27,51 | positive |
|  | 3945 | 32,89 | 33,5 | 27,28 | positive | 32,45 | 37,75 | 27,68 | positive |
|  | 3958 | 33,58 | 33,05 | 25,42 | positive | 35,28 | 37,97 | 28,37 | positive |
|  | **3231** | **33,93** | **40,97** | **27,3** | **undetermined** | **32,24** | **36,81** | **29,4** | **positive** |
|  | 3831 | 34,93 | 40,21 | 27,75 | undetermined | 37,09 | <u>42</u> | 29,76 | undetermined |
|  | 3879 | 35,14 | 36,84 | 25,89 | positive | 33,49 | 37,92 | 26,24 | positive |
|  | 3413 | 35,45 | 38,36 | 26,13 | positive | 26,93 | 30,07 | 26,32 | positive |
|  | **2882\*** | **35,92** | **39,8** | **28,82** | **positive** | **36,73** | **41,97** | **32,17** | **undetermined** |
| Cq N1 > 36 | 3880 | 36,09 | 39,02 | 27,42 | positive | 34,91 | 41,8 | 27,92 | positive |
|  | 3965 | 36,47 | 39,95 | 25,96 | positive | 34,43 | 38,95 | 26,68 | positive |
|  | **3285** | **36,61** | **40,03** | **26** | **undetermined** | **32,06** | **37,47** | **27,89** | **positive** |
|  | 3409 | 38,68 | 39,06 | 25,2 | positive | 31,4 | 34,96 | 25,44 | positive |
|  | 3474 | 36,92 | <u>42</u> | 28,33 | undetermined | 35,93 | 41,71 | 28,14 | undetermined |
|  | 2776* | 37,57 | 38,19 | 27,91 | positive | 30,18 | 36,18 | 27,92 | positive |
|  | **3298** | **37,64** | **42** | **26,22** | **undetermined** | **32,07** | **36,86** | **35,89** | **positive** |
|  | **3197\*** | **37,93** | **41,99** | **25,98** | **undetermined** | **42** | **42** | **25,48** | **negative** |
|  | 2867* | 38,51 | 39,18 | 26,1 | positive | 33,91 | 37,86 | 27,41 | positive |
|  | 3471 | 38,00 | 41,94 | 30,3 | undetermined | 35,72 | 40,73 | 29,21 | undetermined |
|  | 3479 | 38,91 | <u>42</u> | 30,28 | undetermined | 37,22 | 40,97 | 30,24 | undetermined |
|  | **2946\*** | **38,95** | **42,31** | **28,11** | **undetermined** | **42** | **42** | **30,8** | **negative** |
|  | **2815\*** | **39,25** | **41,2** | **31,14** | **undetermined** | **32,52** | **38,67** | **28,32** | **positive** |
|  | **2943\*** | **39,96** | **42** | **26,52** | **undetermined** | **42** | **42** | **24,91** | **negative** |
| Cq N1 >40 | 2517 | <u>42</u> | <u>42</u> | 22,61 | Negative | <u>42</u> | <u>42</u> | 26,55 | negative |
|  | 2518 | <u>42</u> | <u>42</u> | 25,98 | Negative | <u>42</u> | <u>42</u> | 33,63 | negative |
|  | 2927* | <u>42</u> | <u>42</u> | 29,55 | negative | <u>42</u> | <u>42</u> | 27,76 | negative |
|  | 3877 | <u>42</u> | <u>42</u> | 29,21 | negative | <u>42</u> | <u>42</u> | 28,97 | negative |
|  | 3878 | <u>42</u> | <u>42</u> | 27,11 | negative | <u>42</u> | <u>42</u> | 26,31 | negative |
|  | 3881 | <u>42</u> | <u>42</u> | 26,07 | negative | <u>42</u> | <u>42</u> | 25,95 | negative |
|  | 3882 | <u>42</u> | <u>42</u> | 25,13 | negative | <u>42</u> | <u>42</u> | 24,19 | negative |
|  | 3973 | <u>42</u> | <u>42</u> | 29,03 | negative | <u>42</u> | <u>42</u> | 30,24 | negative |
|  | 3960 | <u>42</u> | <u>42</u> | 25,8 | negative | <u>42</u> | <u>42</u> | 24,94 | negative |
|  | 3961 | <u>42</u> | <u>42</u> | 29,96 | negative | <u>42</u> | <u>42</u> | 31,62 | negative |
|  | 3962 | <u>42</u> | <u>42</u> | 29,15 | negative | <u>42</u> | <u>42</u> | 33,11 | negative |
|  | 3963 | <u>42</u> | <u>42</u> | 26,82 | negative | <u>42</u> | <u>42</u> | 28,16 | negative |
|  | 3964 | <u>42</u> | <u>42</u> | 29,18 | negative | <u>42</u> | <u>42</u> | 32,58 | negative |
|  | 4170 | <u>42</u> | <u>42</u> | 29,85 | negative | <u>42</u> | <u>42</u> | 34,6 | negative |
|  | 4173 | <u>42</u> | <u>42</u> | 27,25 | negative | <u>42</u> | <u>42</u> | 32,82 | negative |

|  |  |  |  |  |  |  |  |  |
| --- | --- | --- | --- | --- | --- | --- | --- | --- |
| 4174 | <u>42</u> | <u>42</u> | 28,4 | negative | <u>42</u> | <u>42</u> | 32,92 | negative |
| 4175 | <u>42</u> | <u>42</u> | 29,63 | negative | <u>42</u> | <u>42</u> | 33,72 | negative |

The \* denotes extraction was done with 600 µL of Lysis Buffer. All other samples were extracted using 300 µL as described in Materials and Methods. Bold letters show samples that changed their report's results. A Cq value of 42 was considered for those negative q-PCR results where no Cq value is provided in order to calculate the difference between Cq values.

**Figure 2.**
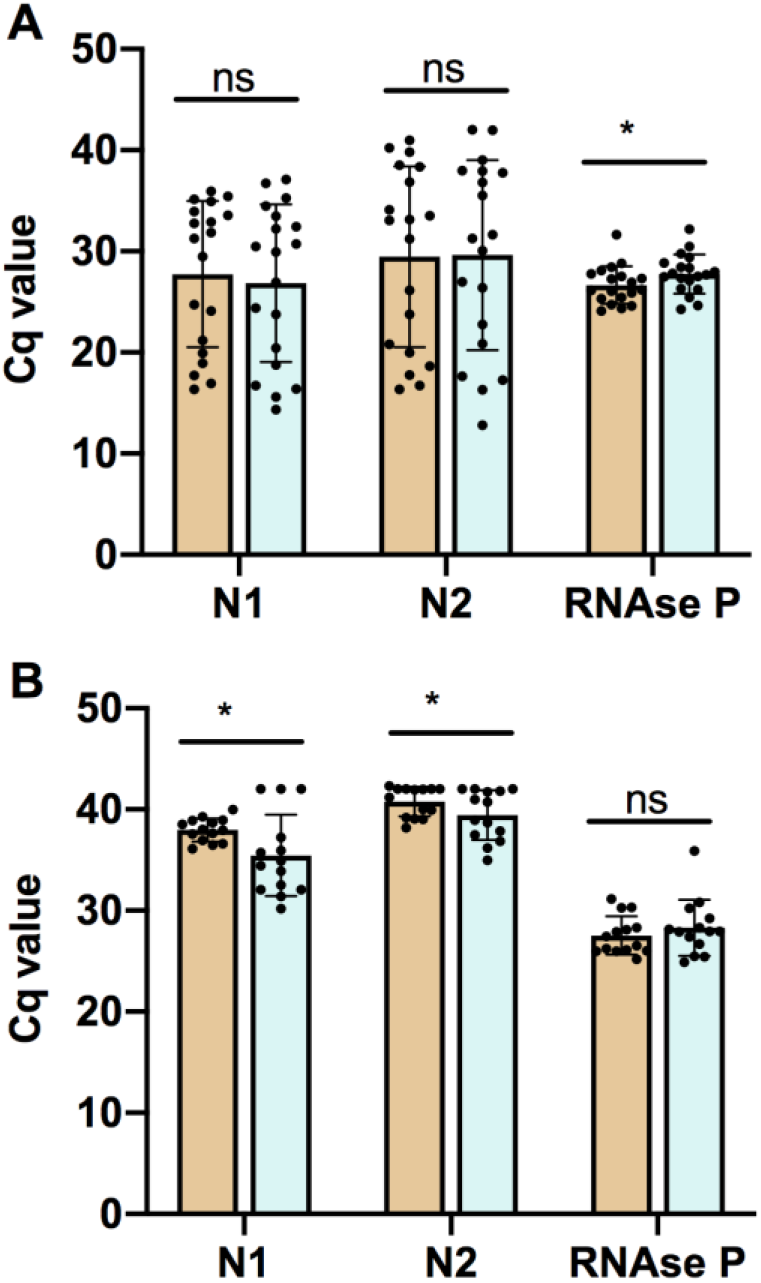
The acid-pH method provides comparable results to commercial kits in clinical samples. Bars represent the mean +/− standard deviation Cq values for each RT-qPCR target gene N1, N2, and RNase P, for samples with Cq N1 ≤ 36 (A) and with Cq N1 >36 (B). Each dot represents one sample. Orange bars show results obtained with High Pure Viral RNA Kit (Roche). Blue bars show results obtained with the acid pH method. Pairwise comparisons of mean Cq values for each target gene were done using a two-tailed p r Stu t’s t-test, with a confidence level of 95% ‘ s’ s statistically significant differences.

Importantly, the processing time and laboriousness of the acid-pH method is similar or less than that of High Pure Viral RNA Kit (Roche) method. A detailed scheme of the method is shown in **Figure 3**.

**Figure 3.**
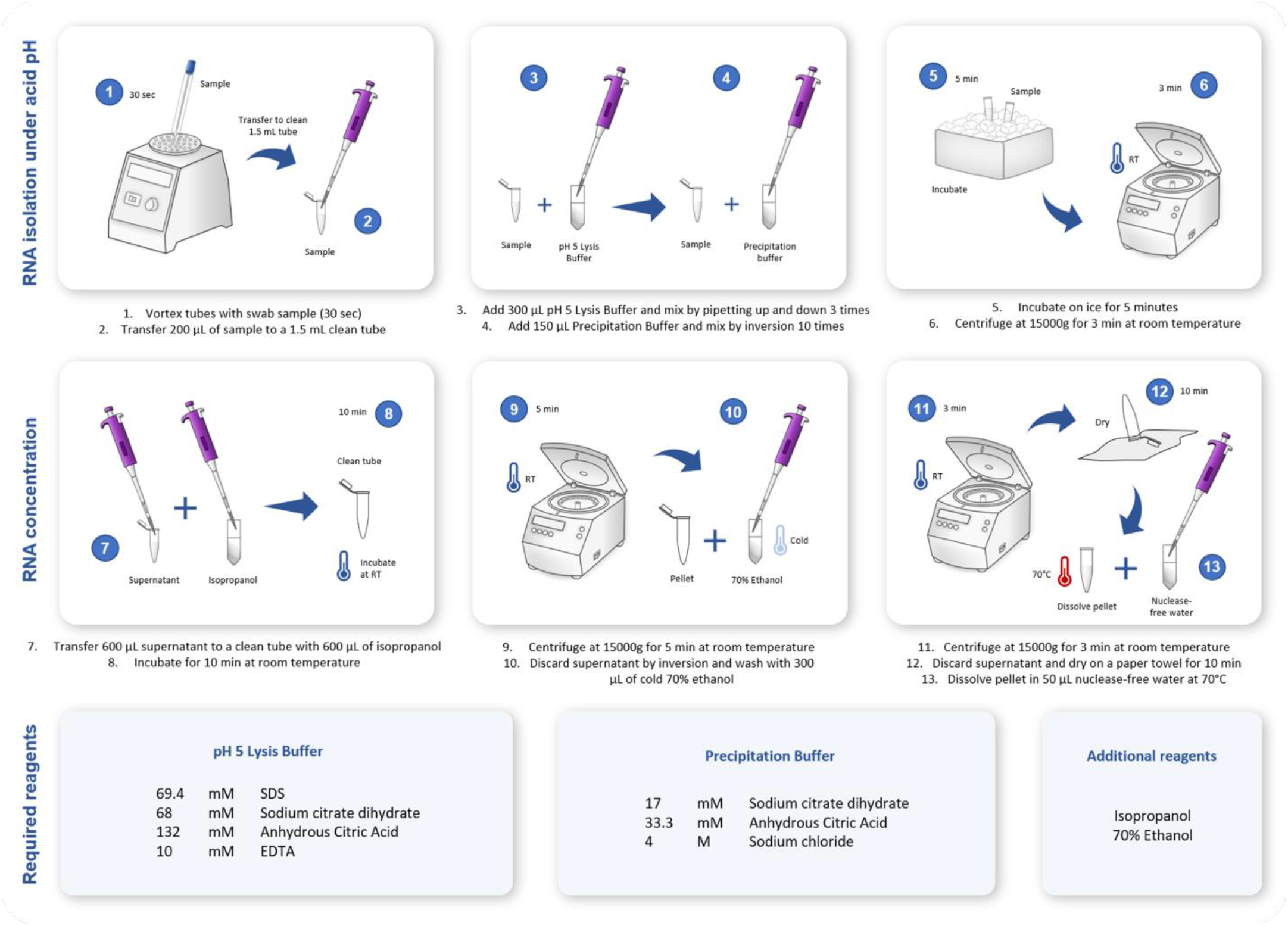
Schematic diagram of the validated acid-pH method for RNA extraction compatible with SARS-CoV2 RT-PCR testing. Steps carried out in the acid pH RNA extraction protocol.

## Discussion

Here we tested several kit-free RNA extraction methods compatible with RT-qPCR analysis and selected one simple procedure based on RNA extraction using acid pH. We validated this method using 50 clinical samples with results comparable to those obtained with commercial kits. There are three key aspects of this method that must be pointed out. First, the acid pH-based methods that we reviewed^12,14,22^ are intended for RNA extraction from tissue, cultured cells, and cell-associated virus. Therefore, the first step of these protocols is centrifugation with subsequent lysis of the cell pellet. However, we need to recover free viral particles in solution, which do not sediment after routine centrifugation at 15,000 g. For this reason we used the uncentrifuged sample directly mixed with lysis buffer, with subsequent precipitation of viral RNA in the whole mix volume. Using uncentrifuged sample is the key step for efficient RNA recovery because when centrifuged sample was used in preliminar tests, Cq values were much higher than those obtained with High Pure Viral RNA Kit (Roche). Second, the acid pH method uses the anionic detergent Sodium dodecyl sulfate (SDS) that can lyse cells and viral coats through disruption of noncovalent bonds in proteins causing them to lose their native conformation^12^. Third, low pH and high concentration of salt make possible the selective recovery of RNA. Within the pH range of 5.5 to 6.0, RNA degradation is minimized^22^. RNA phosphodiester bond is more stable at acidic than alkaline pH, where it is susceptible to alkaline hydrolysis at pH greater than 6^23^. Acid hydrolysis can only occur at pH lower than 2^12,24^. Moreover, DNA and RNA have different solubility at different pH, mainly due to the 2' hydroxyl group of RNA, which increases the polarity of this nucleic acid^25,26^. Therefore, it is essential to adjust the Lysis Buffer to pH 5, as described in Materials and Methods.

It is worth mentioning that all of the samples that changed their report had Cq values that were around the cutoff value of 40. These changes occurred in both directions, meaning that some Cqs increased and some Cqs decreased. It would have been very clarifying to perform triplicated RNA extractions, in particular for undetermined samples, whose viral load is around the detection limit. Because of the above exposed information we consider the acid pH method robust and reliable. In fact, it is currently being used in our diagnostic laboratory since the 3^rd^ week of April 2020 for routine detection of SARS-CoV2 in clinical samples.

The RNA extraction procedure with acid pH described here has many advantages over commercial kits to test for SARS-CoV-2 in the context of the current pandemic. This experimental procedure utilizes low cost reagents and equipment that can be found in standard molecular biology laboratories. The cost of extraction is a critical issue in most clinical laboratories, and the cost of our in-house method is around ten times lower than extraction kits. Moreover, DNase treatment is not necessary because SARS-CoV-2 detection is not altered in the presence of DNA. In fact, residual DNA may serve as the template for RNase P gene amplification. Because of current environmental concerns, we would also like to highlight the lower plastic contamination generated by this in-house method. Column-based extraction kits use several disposable tubes per sample, columns, bottles of buffer solutions, and plastic bags. Our in-house extraction method is by far, much more environmental friendly; it requires only two Eppendorf tubes per sample. Finally, our in-house method is comparable in hands-on time to commercial kits: it can be carried out in approximately 40 minutes for a set of 10 samples. However, it is important to mention that additional care must be taken in handling to avoid cross-contamination between samples.

In conclusion, the RNA extraction procedure with acid pH described here is an excellent alternative to commercial systems to test for SARS-CoV2. Our results support a new method for RNA extraction from swab samples that can be used to detect SARS-CoV2 by standard RT-qPCR testing protocols. This procedure can be a helpful alternative for laboratories facing supply-chain disruption and commercial kit shortages.

## Materials and Methods

### Biological samples

Two types of biological samples were used. For preliminary evaluation of the RNA extraction methods we used saliva samples obtained from two asymptomatic volunteers. Saliva is routinely collected for the initial assessment of viral infection. Two saliva samples were obtained from each volunteer and at least three independent RNA extractions were performed from each sample, obtaining a minimum of six RNA preparations to test each experimental procedure. For validation of the RNA extraction method selected, we used nasopharyngeal swabs in Universal Transport Medium (UTM). Swabs were obtained from 50 patients that attended the outpatient service of Red Salud UC-CHRISTUS (Santiago, Chile) because of suspected coronavirus infection. Only one sample was obtained per patient: one portion of the sample was extracted using the High Pure Viral RNA Kit (Roche), and another portion of the same sample was extracted using the acid pH method. Informed consent was obtained from all participants and/or their legal guardians. Samples were processed in the Laboratory of Diagnostic Microbiology of the same institution. All methods were performed in accordance with the relevant guidelines and regulations. All procedures were approved by the Ethics Committee of the Pontificia Universidad Católica de Chile.

### RNA extraction methods evaluated

The following experimental procedures were tested in this study. Saliva samples were centrifuged before taking an aliquot of supernatant for processing as described below.

#### (1) TRIzol

The standard TRIzol-based method was evaluated^9,11,18^. First, 800 μL of TRIzol were added to 200 μL of sample and vortexed briefly. Then, 200μL of chloroform were added, vortexed, and centrifuged at 12,000 g for 10 min at room temperature. The aqueous phase (600 μL) was recovered in a clean tube containing 600 μL of isopropanol. The tube was mixed by inversion and incubated at room temperature for 10 min. The tube was then centrifuged at 12,000 g for 10 min at 4°C, and the supernatant was discarded. The pellet was washed with 500 μL 70% ethanol, centrifuged at 7,500 g for 5 min at 4°C and the supernatant was discarded. The pellet was dried at room temperature for 10 min and resuspedned in 25 μL RNase-free water by incubating at 37°C for 10 min.

#### (2) BSA-based method

Previous reports show that BSA has positive effects on RT-qPCR results when added to samples in the presence of inhibitors^27,28^. Based on the procedure described by Plante *et al.* (2010)^27^ and Svec *et al.* (2013)^28^, 200 μL aliquot sample was centrifuged at 12,000 for 30 s at room temperature. Then, 2.5 μL of supernatant were added to 47.5 μL of a 1 mg/mL BSA solution (1:20 ratio), vortexed for 30 s and kept on ice or at −80°C until further use.

#### (3) Acid pH-based method

Under acidic pH, RNA can be separated from DNA and other molecules due to the differential polarity given by its hydroxyl groups, which maintains it in solution^12,22,25,26,29^. Based on the methods described by Heath (1999)^22^, Sambrook and Russell (2001)^12^, and Chomczynski and Sacchi (2006)^14^, 300 μL of pH 5 Lysis Buffer (69,4 mM SDS, 68 mM sodium citrate dihydrate, 132 mM anhydrous citric acid and 10 mM EDTA, then adjust the buffer to pH 5) were added to 200 μL of uncentrifuged sample and mixed by pipetting three times. Then, 150 μL of Precipitation Buffer (17 mM sodium citrate dihydrate, 33,3 mM anhydrous citric acid, and 4 M NaCl) were added and mixed by inversion 10 times. Samples were incubated on ice for 5 min and centrifuged at 15,000 g for 3 min at room temperature. 600 μL of the supernatant were transfered to a clean tube containing 600 μL s pr p ub t r 10 min at room temperature. A new centrifugation step was made at 15,000 g for 5 min at room temperature. The supernatant was removed, and the pellet was w t 300 μL cold 70% ethanol and centrifuged at 15,000 g for 3 min at room temperature. Supernatant was discarded and tubes were inverted in paper towel. The pellet was dried, leaving the tubes open for 10 min. Finally, the pellet was resuspended i 50 μL nuclease-free water pre-warmed at 70°C.

NOTE: If the buffer is stored for later use, it precipitates at 4°C, so it needs to be heated for 5 minutes at 60°C for its use.

#### (4) High temperature-based method

Based on the method described by Fomsgaard and Rosenstierne (2020)^30^, 50 μL of the sample were directly heated at 98°C for 5 min and cooled at 4°C. Then 19 μL of the sample were mixed with 1 μL of BSA (20 mg/mL) and kept on ice for immediate use or at −80 °C for later use.

#### (5) Direct use of the samples

An aliquot taken from the original sample was directly used to perform RT-qPCR analysis^31^.

The 50 nasopharyngeal swabs used for the validation of the RNA extraction method selected, were extracted using High Pure Viral RNA Kit (Roche) according to instructions provided by the manufacturer. This RNA extraction method was considered as the gold standard for comparison purposes, and It is based in capture of RNA using columns with silica filters.

### RT-qPCR analysis

For preliminary evaluation of RNA extraction procedures, we used RT-qPCR against the human RNAse P gene with primers and a Taqman probe previously described^32^. RP1-F: AGATTTGGACCTGCGAGCG, RP1-R: GAGCGGCTGTCTCCACAAGT, and RP1-probe: TTCTGACCTGAAGGCTCTGCGCG. The RNase P gene is used as an internal control because many copies of it exist in the human genome, and it is readily detectable. The source of RNase P comes from the human cells that are present in every sample used. It is assumed that if human nucleic acids were extracted to detect the human gene RNase P, viral nucleic acids were also successfully extracted. The RNase P target is also amplified as a quality control for the extraction method and to corroborate the absence of PCR-inhibitors in the sample.

For RT-qPCR 5 μL of RNA from saliva samples, 2 μL of RNase-Free water 1 μL of each RNase P prime, 1 μL of TaqMan RNase P probe and 10 μL of 2X TaqMan Fast Universal PCR Master Mix (Applied Biosystems), were used in a final reaction volume of 20 μL performed in a StepOnePlus Real-Time PCR System (Applied Biosystems).

For validation of the selected RNA extraction procedure, RT-qPCR using Taqman probes and primers recommended by the CDC was used^21^. Two viral targets were amplified: the nucleocapsid viral proteins N1 and N2. Primers and probe for N1 were N1-F: GACCCCAAAATCAGCGAAAT, N1-R: TCTGGTTACTGCCAGTTGAATCTG, and N1-probe: FAM-ACCCCGCATTACGTTTGGTGGACC-BHQ1. Primers and probe for N2 were N2-F: TTACAAACATTGGCCGCAAA, N2-R: GCGCGACATTCCGAAGAA, and N2-probe: FAM-ACAATTTGCCCCCAGCGCTTCAG-BHQ1. Primers and probe for RNase P were RP2-F: AGATTTGGACCTGCGAGCG, RP2-R: GAGCGGCTGTCTCCACAAGT, and RP2-probe: FAM–TTCTGACCTGAAGGCTCTGCGCG–BHQ1. A one-step RT-qPCR reaction was performed in a StepOnePlus Real-Time PCR System (Applied Biosystems). Cutoff points for Cq values (Cycle of quantification, or Cycle Threshold) required to decide whether a result is COVID-19 positive or negative were those specified by CDC as follows. To report a positive result, both viral targets N1 and N2 must be Cq<40. To report a negative result both viral targets must be Cq≥40. If one of the viral targets is Cq<40 and the other is Cq≥40 the result must be reported as undetermined. The RNase P target must be Cq≤35

### Statistical analysis

Mean Cq values obtained through both methods for each target gene were analyzed in pairwise comparisons using a p r Stu t’s t-test. The analysis was performed using GraphPad Prism 8 software.

## Acknowledgments

We would like to acknowledge people that contributed with helpful discussions and critical comments that helped us along the way, especially Professor Francisco Melo and Professor Marcelo López-Lastra. We would like to also thank Maite Salazar and Laura Delgado for helping with language edits. We would like to express our gratefulness to technicians of the Laboratory of Diagnostic Microbiology, Sandra Prado and Javier Hernández, for all their help with technicals aspects of this work.

## Author Contributions

AW performed experimental work, data analysis, figure composition, and manuscript writing. AC, CIH, and VS did experimental work, data analysis, and figure composition. GA, LL, performed a literature search and manuscript writing. CM, ML, SS, AMG, and TQ contributed to experimental design and critical data analysis. SH, ER, MF, performed a literature search and defined protocols for testing. RAG and PG performed the literature search, experimental design, data analysis, and manuscript writing. They co-supervised this project. All authors reviewed the manuscript.

## Additional Information

The authors declare no competing interests.

